# Decomposition of the SARS-CoV-2-ACE2 interface reveals a common trend among emerging viral variants

**DOI:** 10.1101/2021.05.28.446149

**Authors:** Eileen Socher, Marcus Conrad, Lukas Heger, Friedrich Paulsen, Heinrich Sticht, Friederike Zunke, Philipp Arnold

## Abstract

New viral variants of the SARS-CoV-2 virus show enhanced infectivity compared to wild type, resulting in an altered pandemic situation in affected areas. These variants are the B.1.1.7 (United Kingdom), B.1.1.7 with the additional E484K mutation, the B.1.351 variant (South Africa) and the P.1 variant (Brazil). Understanding the binding modalities between these viral variants and the host cell receptor ACE2 allows depicting changes, but also common motifs of virus-host cell interaction. The trimeric spike protein expressed at the viral surface contains the receptor-binding domain (RBD) that forms the molecular interface with ACE2. All the above-mentioned variants carry between one and three amino acid exchanges within the interface-forming region of the RBD, thereby altering the binding interface with ACE2. Using molecular dynamics simulations and decomposition of the interaction energies between the RBD and ACE2, we identified phenylalanine 486, glutamine 498, threonine 500 and tyrosine 505 as important interface-forming residues across viral variants. We also suggest a reduced binding energy between RBD and ACE2 in viral variants with higher infectivity, attributed to residue-specific differences in electrostatic interaction energy. Importantly, individual amino acid exchanges not only influence the affected position, but also alter the conformation of surrounding residues and affect their interaction potential as well. We demonstrate how computational methods can help to identify changed as well as common motifs across viral variants. These identified motifs might play a crucial role, in the strategical development of therapeutic interventions against the fast mutating SARS-CoV-2 virus.

**Significance Statement:** The COVID-19 pandemic caused by the SARS-CoV-2 virus has significantly changed our lives. To date, there is a lack of neutralizing drugs that specifically target SARS-CoV-2. Hope lies in newly developed vaccines that effectively prevent severe cases of acute respiratory syndrome. However, emerging viral variants escape vaccine-induced immune-protection. Therefore, identification of appropriate molecular targets across viral variants is important for the development of second- and third-generation vaccines and inhibitory antibodies. In this study, we identify residues across viral variants that are important for viral binding to the host cell. As such residues cannot be replaced without diminishing infectivity of the virus, these residues represent primary targets for intervention, for example by neutralizing antibodies.

## Introduction

The COVID19 pandemic caused by the SARS-CoV-2 virus is having a major impact on human lives worldwide (1, 2). For cellular infection, the virus engages the cell surface protein angiotensin converting enzyme 2 (ACE2) via its trimeric spike protein (3, 4). Within the spike protein, the receptor-binding domain (RBD) interfaces with ACE2 (5, 6) (Fig. 1A), mediating binding of the virus to the host cell surface. Priming at the S2’ cleavage site within the spike protein by the serine protease TMPRSS2 releases the fusion peptide from the protein backbone (3, 7, 8). After insertion of the fusion peptide into the host cell membrane and the activation of a conformational switch leading to dissociation of the spike protein’s S1 and S2 domains (9–11), the virus and host cell membrane come close and fuse (7, 12). To allow the two membranes to approach, all non-cleaved subunits of the spike protein must dissociate from the ACE2 receptors, to avoid steric hindrance (5, 13, 14). Thus, optimized binding efficiency between the RBD and ACE2 is important for viral entry (14). In several viral variants that showed superior infectivity over the wild type (wt) SARS-CoV-2 virus, amino acid exchanges at the interface of the RBD and ACE2 have been reported (15, 16). In this study, we examined the RBD variants of B.1.1.7 (originated in the UK), B.1.1.7+E484K (also UK), B.1.351 (South Africa) and P.1 (Brazil) in complex with ACE2 using molecular dynamics (MD) simulations. The B.1.1.7 variant accommodates an N501Y exchange within the RBD-ACE2 interface and the B.1.1.7+E484K exhibits an additional glutamate-to-lysine exchange at position 484 (15). The B.1.351 and P.1 variants carry both of these mutations and have additional exchanges at amino acid position 417: in B.1.351, the lysine at position 417 is exchanged for an asparagine (K417N) and in P.1 for a threonine (K417T) (15). Notably, the exchange of lysine for glutamate at position 484 is associated with a reduced efficacy for certain vaccines (17, 18). In this study, we demonstrate that progressive viral variants occur with reduced binding energies to ACE2. Additionally, we show how decomposition of the RBD-ACE2 interface can identify residues on the RBD that are critical for virus host interactions across different SARS-CoV-2 variants. Analyses of electrostatic and van der Waals linear interaction energies reveal the amino acids phenylalanine 486, glutamine 493, and tyrosine 505 as critical residues for binding of the RBD to ACE2 across all RBD variants examined. Our data helps to identify epitopes that are common in all viral variants and might serve as targets for the development of neutralizing antibodies.

**Figure 1.**
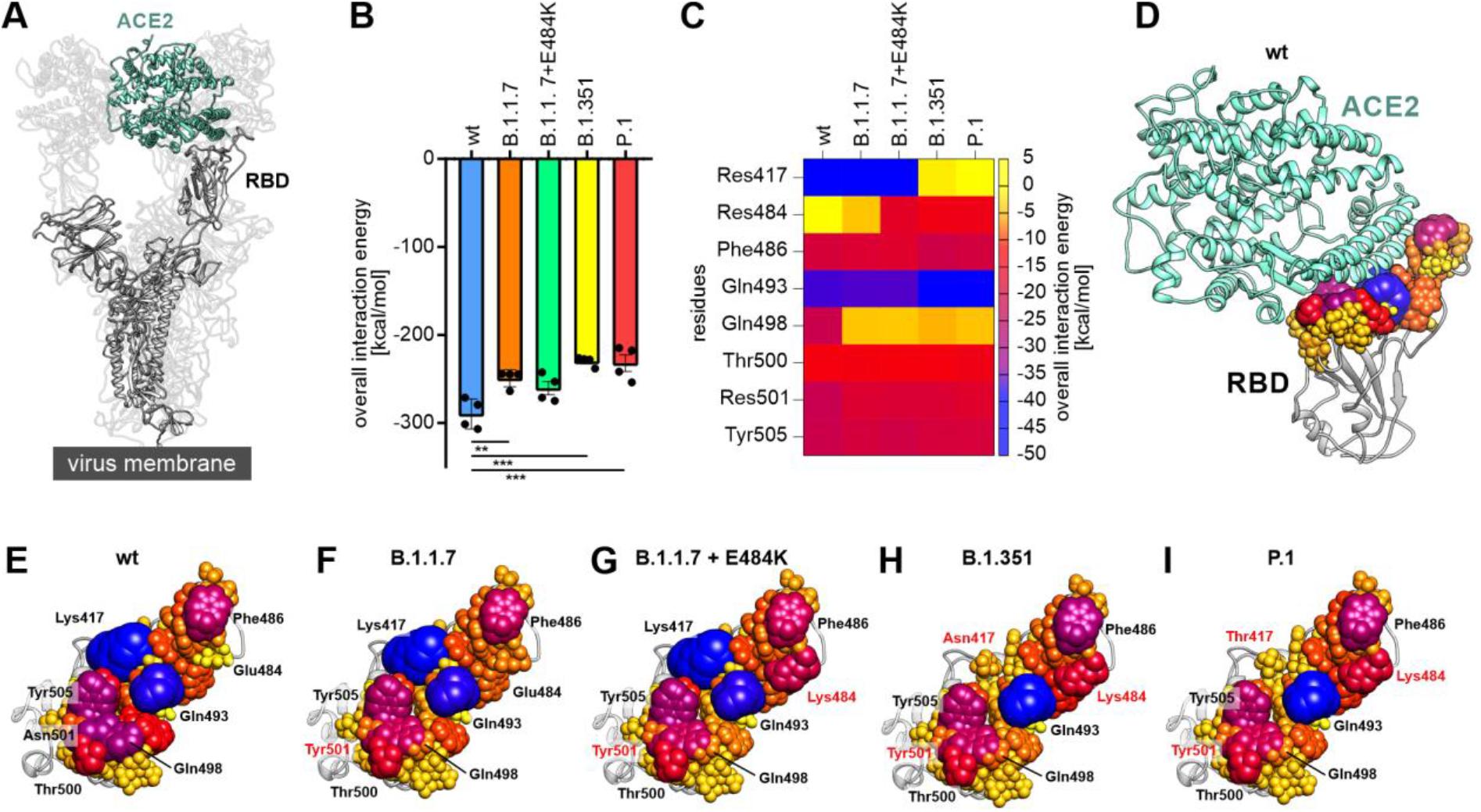
Decomposition of the overall linear interaction energies. (A) Structural representation of the trimeric spike protein as expressed on the viral membrane (grey) with exposed receptor binding domain (RBD). The host cell receptor ACE2 is displayed in aquamarine (PDB ID: 7kms (19)). (B) Overall linear interaction energy as calculated for the different spike protein variants (wt = wild type). Statistical analysis was performed using one-way ANOVA (n=4, differences assumed significant for ** p<0.01, *** p<0.001). (C) Heat map of receptor-binding domain (RBD) expressed residues important for binding of ACE2. (D) Wild type RBD in complex with ACE2 (aquamarine). All residues within a maximum distance of 8 Å to ACE2 are displayed according to their interaction energy with different colors and sphere radii. (E) View on the interface formed by the wild type RBD with residues displayed in different colors and sphere diameters according to their interaction energy. Especially lysine 417 (Lys417), glutamate 484 (Glu484), phenylalanine 486 (Phe486), glutamine 493 (Gln493), glutamine 498 (Gln498), threonine 500 (Thr500), asparagine 501 (Asn501) and tyrosine 505 (Tyr505) were of interest. (F) View on the interface formed by the RBD of the B.1.1.7 variant with ACE2. Note that residue 501 is mutated to tyrosine (Tyr501; red) in this variant. (G) View on the interface formed by the RBD of the B.1.1.7+E484K variant with ACE2. Note the additional change of residue 484 from glutamate to lysine (Lys484; red) compared to wild type and B.1.1.7. (H) View on the interface formed by the RBD of the B.1.351 variant with ACE2. This variant carries the tyrosine at position 501, the lysine at position 484 and an additional exchange from lysine to asparagine at position 417 (Asn417; red). (I) View on the interface formed by the RBD of the P.1 variant with ACE2. Besides a tyrosine at position 501 and a lysine at position 484 this variant carries a lysine to threonine exchange at position 417 (Thr417; red).

## Results

### Mutations reduce and balance the interaction energy between RBD and ACE2

To calculate the interaction energy between the cell surface receptor ACE2 and different RBD variants, the structure of the wild type RBD and ACE2 complex (PDB ID: 7kmb (19)) was used as starting point. Variant-specific amino acid substitutions were introduced in the RBD sequence before applying MD simulations: B.1.1.7 (N501Y), B.1.1.7+E484K (N501Y, E484K), B.1.351 (N501Y, E484K, K417N) and P.1 (N501Y, E484K, K417T). After simulation for 500 ns, we calculated the overall linear interaction energy between the different RBDs and the host cell receptor ACE2 (Fig. 1B). We found the strongest interaction of ACE2 with the wild type, which was significantly stronger compared to variants B.1.1.7, B.1.351 and P.1. We also calculated the binding energy using molecular mechanics/Generalized Born surface area (MM/GBSA) and found a similar trend (Fig. S1A). When we plotted the sum of the electrostatic and the van der Waals parts of the linear interaction energy for all RBD residues within 8 Å to ACE2 (Fig. S1B), we identified eight residues that we analyzed in more detail (Fig. 1C): amino acid positions 417 (lysine, asparagine, or threonine), 484 (glutamate or lysine), 486 (phenylalanine), 493 (glutamine), 500 (threonine), 501 (asparagine or tyrosine), and 505 (tyrosine). The distribution of the overall interaction energy from the RBD in complex with ACE2 revealed the central part of the RBD as the main contact site (Fig. 1D for wild type; Fig. S1C for B.1.1.7, B.1.1.7+E484K, B.1.351 and P.1). Illustrating the interaction interface of the wild type RBD with all residues as spheres with radii and colors according to their overall linear interaction energy (yellow (5 kcal/mol) – blue (−50 kcal/mol)), demonstrated that lysine 417 and glutamine 493 are major interaction partners with ACE2 (Fig. 1E). Both amino acids are flanked by phenylalanine 486 on one side and glutamine 498, threonine 500, asparagine 501, and tyrosine 505 on the other side (Fig. 1E). Depiction of the mutant RBD binding interfaces revealed significant structural changes within the ACE2 interaction site. For example, B.1.1.7 showed a major change in the RBD as glutamine 498 loses its interaction energy compared with the wild type variant (Fig. 1E, F). In the B.1.1.7+E484K variant, the newly inserted lysine at position 484 introduces interaction energy in an area close to phenylalanine 486 (Fig. 1G). Exchange of lysine 417 to asparagine (B.1.351, Fig. 1H) or threonine (P.1, Fig. 1I) reduced the importance of amino acid 417 in ACE2 binding. This resulted in glutamine 493 as the most important interacting residue flanked by two balanced interaction sites formed by phenylalanine 486 and lysine 484 on one side and threonine 500, tyrosine 501, and tyrosine 505 on the other side (Fig. 1H, I). We also evaluated the root mean square fluctuation (RMSF) values for all RBD variants in complex with ACE2 to assess the overall structural flexibility of the different RBD variants, but we found no significant differences in structural dynamics (Fig. S1D).

### Decomposition of the binding interface reveals global changes in electrostatic interaction energy

Analyses of the electrostatic interaction energies between the RBD and ACE2 showed reduced energies for all mutated variants, with significant differences for the B.1.1.7, B.1.351, and P.1 variants compared to wild type (Fig. 2A). Decomposition of the electrostatic linear interaction energy to the level of individual amino acids (Fig. S2A, B) revealed residues 417, 484, 493, and 498 as strongest interaction sites for different spike variants (Fig. 2B). In contrast, the van der Waals linear interaction energy was similar between the different RBD variants analyzed (Fig. 2C), and the decomposition revealed only minor differences for individual residues (Fig. 2D, Fig. S2C, D). To analyze the consequences of individual residues on ACE2 binding at the molecular level, we measured interatomic distances, calculated the number of contacts and linear interaction energies for specific amino acids. Interestingly, we discovered that the lysine at position 417, found in wild type and RBD variants B.1.1.7 and B.1.1.7+E484K, formed a salt bridge with aspartate 30 of the ACE2 receptor (Fig. 2E). Accordingly, the distance plot of the two amino acids lysine 417 (RBD wt) and aspartate 30 (ACE2) showed that both residues were in close proximity (<4 Å) underlining their tight interaction as characteristic for salt bridges (Fig. 2F). This indicates the importance of the amino acid at position 417 in RBD-ACE2 interaction. In the following, we therefore analyzed the effect of mutations at this position as reported for variants B.1.351 and P.1. Calculation of the average numbers of contacts per frame showed that a lysine at position 417 (wt, B.1.1.7, B.1.1.7+E484K) induces ~35 contacts, an asparagine (P.1.351) ~5 contacts and a threonine (P.1) ~2 contacts (Fig. 2G). This is reflected by a strongly reduced electrostatic interaction energy with ACE2 for this residue when comparing RBD variants carrying a lysine (wt, B.1.1.7, B.1.1.7+E484K) or lacking a lysine residue (Asn/Thr; P.1.351, P.1) at position 417 (Fig. 2h). An additional effect that we noticed was the increase in electrostatic interaction energy for glutamine 493 in cases where lysine was absent at position 417, as found for variants B.1.351 and P.1 (Fig. 2I). Since all variants lacking a lysine at position 417 also exhibit an exchange of glutamate to lysine at position 484 (E484K), we assume a compensatory effect counteracting the reduced binding energy.

**Figure 2.**
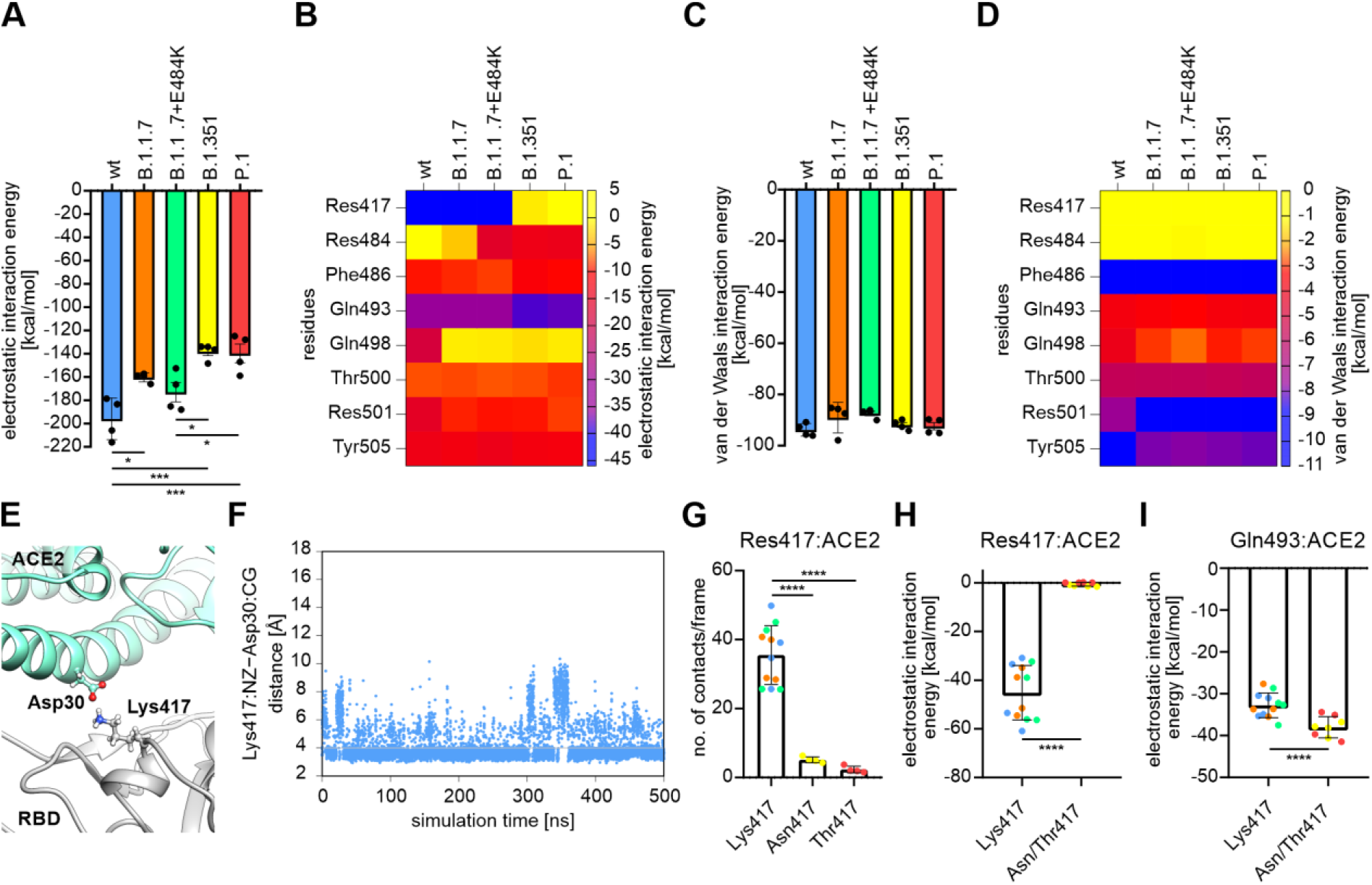
Decomposition into electrostatic and van der Waals linear interaction energies and consequences for spike protein variants exchanging lysine 417. (A) Electrostatic interaction energy as calculated for the different receptor-binding domain (RBD) variants (wt = wild type). Statistical analysis was performed using one-way ANOVA (n=4, differences assumed significant for * p<0.05, *** p<0.001). (B) Heat map of the RBD expressed residues important for ACE2 interaction. Color according to the electrostatic linear interaction energy calculated. (C) Van der Waals linear interaction energy as calculated for the different variants of the RBD. (D) Heat map of the RBD expressed residues important for ACE2 interaction. Color according to the van der Waals linear interaction energy calculated. (E) Structural representation of the salt bridge formed by lysine 417 from the RBD (Lys417) and aspartate 30 from ACE2 (Asp30). (F) Exemplary distance plot, for a simulation from the wild type RBD with ACE2 over time, for lysine 417 and aspartate 30. The grey line indicates a distance of 4 Å. (G) Average number of contacts per frame for residue 417. In variants that express a lysine at this position (wt: blue, B.1.1.7: orange and B.1.1.7+E484K: green) this number is significantly higher when compared to variants with an asparagine (B.1.351: yellow) or threonine (P.1: red; one-way ANOVA; n=4-12; **** p<0.0001). (H) Electrostatic linear interaction energy compared between RBD variants with a lysine (wt: blue, B.1.1.7: orange and B.1.1.7+E484K: green) or asparagine/threonine (B.1.351: yellow and P.1: red; Student’s two-tailed t-test; **** p<0.0001). (I) Electrostatic linear interaction energy of glutamine 493 compared for variants with a lysine (wt: blue, B.1.1.7: orange and B.1.1.7+E484K: green) or an asparagine/threonine (B.1.351: yellow and P.1: red) at position 417 (Student’s two-tailed t-test; *** p<0.001).

### A lysine at position 484 forms a partial salt bridge with glutamate 75 from ACE2

As indicated by mutation and interaction energy data, residue 484 plays a critical role in the RBD-ACE2 binding properties. The exchange from glutamate to lysine at position 484 (as found in B.1.1.7+E484K, B.1.351 and P.1) changes a negatively charged residue to a positive one, allowing salt bridge formation with negatively charged residues in adjacent ACE2 regions. In this region, we identified glutamate 75 expressed on ACE2 as a potential interaction partner (Fig. 3A). Plotting the total fraction of contacts for residue 484 (as glutamate or lysine) with residues expressed on ACE2 revealed increased interaction with glutamate 75 for variants expressing a lysine at position 484 (B.1.1.7+E484K, B.1.351 and P.1) (Fig. 3B). All variants showed interaction with leucine 79. Comparison of variants expressing a glutamate (wt and B.1.1.7) or a lysine (B.1.1.7+E484K, B.1.351 and P.1) at position 484 also showed an increase in the total number of contacts per frame (Fig. 3C). Comparison of the electrostatic linear interaction energies for variants expressing glutamate or lysine at position 484 demonstrated a significantly increased interaction energy for lysine-expressing variants. Thus, our analysis supports the idea that the exchange of a lysine at position 484 for a glutamate increases the interaction energy and thereby stabilizes the RBD-ACE2 interaction. It also introduces electrostatic interaction in close proximity to the important van der Waals interaction formed by phenylalanine 486 (Fig. 1G-I).

**Figure 3.**
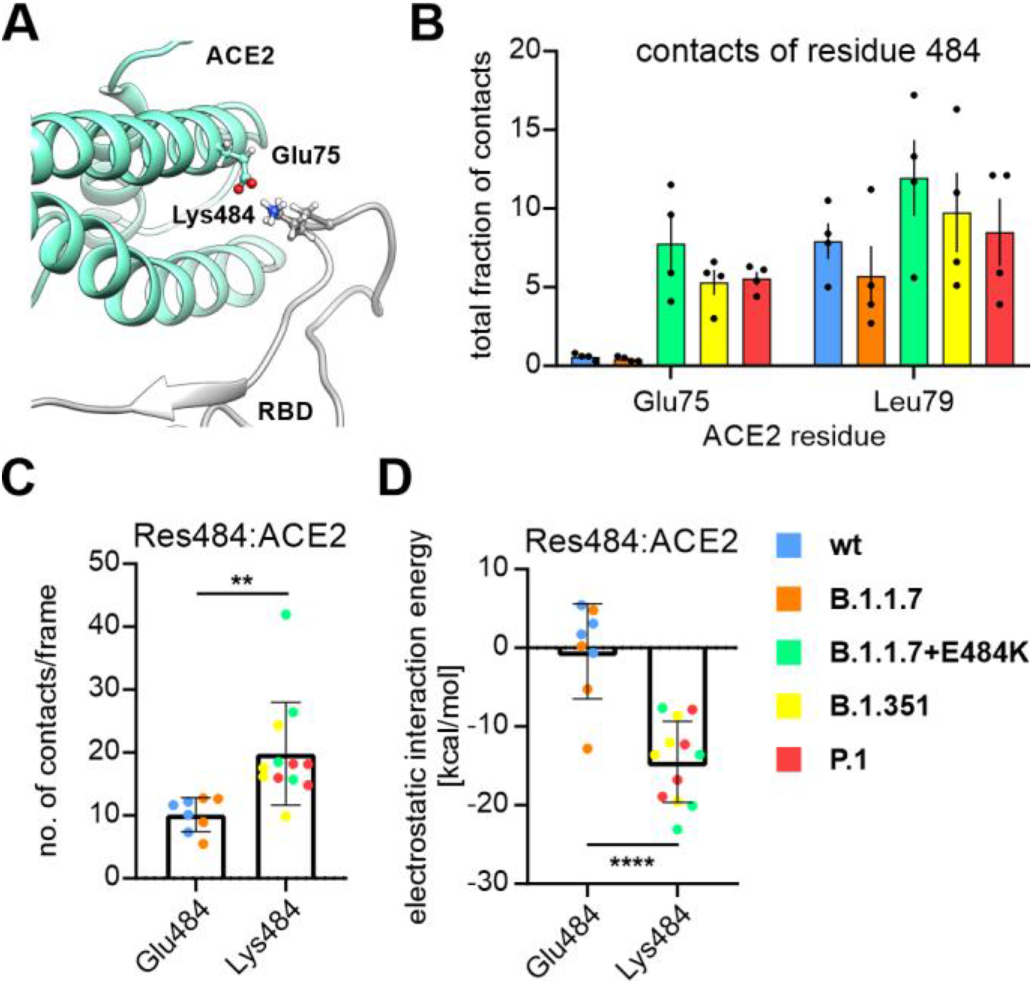
Understanding the changes induced by the E484K mutation in terms of RBD-ACE2 contact formation and increase in local interaction energy. (A) Structural representation of lysine 484 (Lys484) as expressed on the RBD of B.1.1.7+E484K, B.1.351 and P.1, and glutamate 75 (Glu75) expressed on ACE2. (B) Total fraction of contacts for residue 484 (glutamate or lysine) with glutamate 75 (Glu75) or leucine 79 (Leu79) expressed on ACE2. (C) Number of contacts per frame for residue 484. The number is significantly higher for variants that express a lysine (B.1.1.7+E484K: green, B.1.351: yellow and P.1: red) compared to those expressing a glutamate (wt: blue and B.1.1.7: orange; Student’s two-tailed t-test; ** p<0.01). (D) Electrostatic linear interaction energy compared between variants that express a lysine (B.1.1.7+E484K: green, B.1.351: yellow and P.1: red) or a glutamate (wt: blue and B.1.1.7: orange) at position 484 (Student’s two-tailed t-test; **** p<0.0001).

### The N501Y mutation induces a complex reorganization of local interaction energies

Variants expressing a tyrosine at position 501 instead of an asparagine (all except for wild type) showed a significant decrease in the electrostatic interaction energy between the residue at position 501 and ACE2 (Fig. 4A). Although the van der Waals interaction energy increased for these variants compared to wild type (Fig. 4B), the resulting overall interaction energy for this residue decreased when asparagine was exchanged for tyrosine (Fig. 4C). Analysis of the interaction energy of residues in the vicinity of residue 501 revealed that glutamine 498 lost almost all of its interaction energy (from about −22 kcal/mol to −4 kcal/mol) when a tyrosine was present at position 501 (Fig. 1C). This loss of overall interaction energy can be attributed to a loss of electrostatic interaction energy. Comparative analyses of variants with identical residues at this position (asparagine for wild type and tyrosine for all other variants), revealed a reduced electrostatic interaction energy from about −18 kcal/mol to −2 kcal/mol (Fig. 4D). Analysis of the RBD-ACE2 interface around residue 501, which includes glutamine 498, showed a conformational change for this glutamine induced by the insertion of the tyrosine at position 501 (Fig. 4E, F). Analysis of residues interacting with glutamine 498 on ACE2 revealed major interactions with aspartate 38, tyrosine 41, glutamine 42, leucine 45, and lysine 353 (Fig. 4E, G). In variants expressing a tyrosine at position 501, glutamine lost its contacts completely to aspartate 38 and lysine 353, and partially to tyrosine 41. This can be explained by the bulky tyrosine inserted for an asparagine displacing glutamine 498 from the binding interface. With these results, we demonstrate that single amino acid changes also affect surrounding residues and alter the conformation in their immediate vicinity.

**Figure 4.**
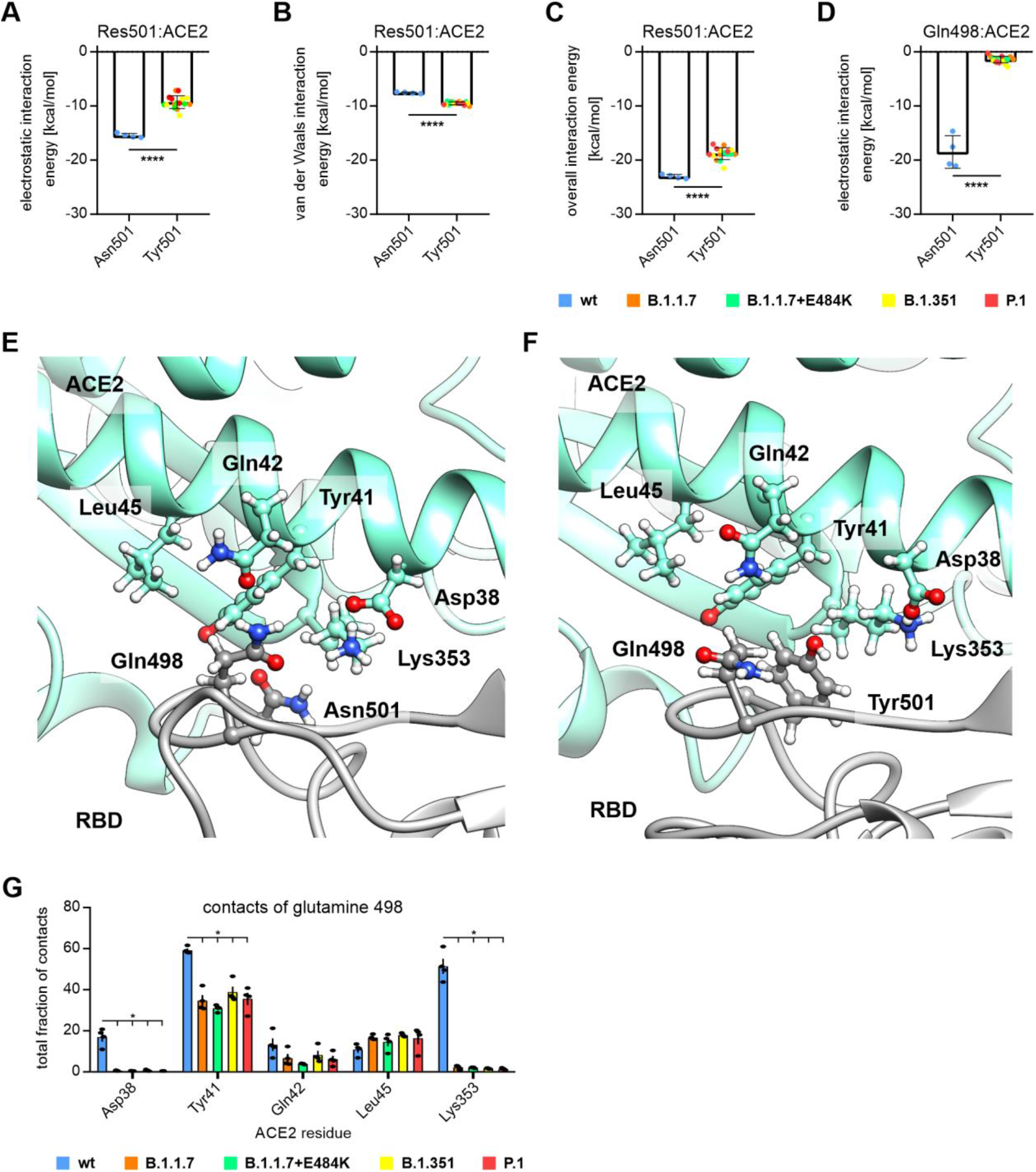
Analysis of the local conformational changes induced by the N501Y mutation. (A) Electrostatic linear interaction energy for residue 501 compared for asparagine expressing wild type (wt: blue) and the tyrosine expressing variants (B.1.1.7: orange and B.1.1.7+E484K: green, B.1.351: yellow and P.1: red). Insertion of a tyrosine decreased the electrostatic interaction (Student’s two-tailed t-test; **** p<0.0001). (B) Van der Waals linear interaction energy increased for variants that express a tyrosine at position 501 (B.1.1.7: orange and B.1.1.7+E484K: green, B.1.351: yellow and P.1: red) when compared to the original asparagine expressing wild type (wt: blue; Student’s two-tailed t-test; **** p<0.0001). (C) Overall interaction energies decreased for variants that express a tyrosine at position 501 (B.1.1.7: orange and B.1.1.7+E484K: green, B.1.351: yellow and P.1: red) when compared to wild type (wt: blue) (Student’s two-tailed t-test; **** p<0.0001). (D) Electrostatic linear interaction energy for glutamine 498 was almost lost for variants that express a tyrosine at position 501 (B.1.1.7: orange and B.1.1.7+E484K: green, B.1.351: yellow and P.1: red) when compared to wild type (wt: blue; Student’s two-tailed t-test; **** p<0.0001). (E) Structural representation of an asparagine 501 expressing wild type receptor-binding domain (RBD) in complex with ACE2. The ACE2 residues aspartate 38 (Asp38), tyrosine 41 (Tyr41), glutamine 42 (Gln42) and leucine 45 (Leu45) were potential interactors for glutamine 498 (Gln498). Gln498 was in close proximity of RBD asparagine 501. (F) Structural representation of a tyrosine 501 expressing variant of the receptor-binding domain (RBD) in complex with ACE2. ACE2 residues aspartate 38 (Asp38), tyrosine 41 (Tyr41), glutamine 42 (Gln42) and leucine 45 (Leu45) present potential interactors for glutamine 498 (Gln498). The bulky tyrosine 501 seemed to partially exclude glutamine 498 from the RBD-ACE2 interface. (G) Total fraction of contacts for glutamine 498 with residues expressed on ACE2. Contacts were mainly lost towards aspartate 38 (Asp38), tyrosine 41 (Tyr41) and lysine 353 (Lys353) in variants that express a tyrosine at position 501 (B.1.1.7: orange and B.1.1.7+E484K: green, B.1.351: yellow and P.1: red; two-way ANOVA; * p < 0.05; full statistical analysis in supplemental table 1).

## Discussion

Understanding the interaction between SARS-CoV-2 and the host cell is important to develop therapeutic antibodies that work across different viral variants. Recently emerged viral variants, such as B.1.1.7, B.1.1.7+E484K, B.1.351, and P.1 display mutations within the RBD at the interface with ACE2 (15, 20). Here we demonstrate how MD simulations allows us to track these changes in interaction energies between RBD and ACE2. A general trend toward reduced interaction energies between the RBD and ACE2 indicated that not only binding, but also subsequent dissociation of the virus from the cell surface receptor serves as a driving force in viral evolution (21, 22). Decomposition of the interaction energies into electrostatic and van der Waals interactions demonstrates that the decrease in electrostatic interaction is induced primarily by the exchange of asparagine 501 for tyrosine and the exchange of lysine 417 to asparagine or threonine. For both mutations at position 417, reduced binding affinity to ACE2 but increased expression of the RBD was measured in a yeast system (23). The E484K mutation induces a local increase in electrostatic interaction energy adjacent to phenylalanine 486. Comparing the spatial distribution of the overall interaction energies for the different variants, it appears that the insertion of N501Y, K417N/T, and E484K as present in B.1.351 and P.1 equalizes the interaction energy uniformly across over the ACE2 binding interface. For the N501Y variants, we can also describe local conformational changes that partially exclude glutamine 498 from its interaction partners on ACE2. These local conformational changes may also play a role in reduced binding of therapeutic antibodies or inadequate protection after vaccination (18, 22, 24).

Our decomposition analysis also allows us to point to residues that seem to have a critical function for RBD-ACE2 interaction across different spike protein variants. These residues are phenylalanine 486, glutamine 493, threonine 500, asparagine/tyrosine501, and tyrosine 505. Further combinatorial mutation experiments on these residues will determine whether they are dispensable for ACE2 binding. From our data, we hypothesize that a loss of glutamine 493 may prevent sufficient binding and thus viral entry. As no spike variant with higher infectious potential has yet been reported to replace glutamine 493, this residue might be a primary target for a neutralizing antibody acting across all SARS-CoV-2 variants. Our data and data from other groups indicate that MD simulation is a powerful tool to evaluate the consequences of individual amino acid exchanges on the interaction energies between the RBD and ACE2 and beyond (25, 26). In particular, for variants with multiple changes or rapidly mutating strains, MD simulations serve as a rapid and powerful tool to understand the impact of each mutation and unravel its mechanistic properties. Validation of these results in biochemical or cell biological experiments and circling back into the simulation enables rapid identification of suitable target sites that can be used for translational approaches.

## Materials and Methods

### Generation of the starting structures

To investigate the interface between the RBD of the spike protein and ACE2, the respective wild type start structure was taken from the PDB database (PDB ID code: 7KMB (19)). To also generate the starting structures for the MD simulations of the different spike variants, the amino acid substitutions (B.1.1.7: N501Y; B.1.1.7+E484K: E484K and N501Y; B.1.351: K417N, E484K and N501Y; P.1: K417T, E484K and N501Y) were introduced with Swiss-PdbViewer 4.1.0.

### Molecular dynamics simulations

Molecular dynamics simulations were performed using version 20 of the Amber Molecular Dynamics software package (ambermd.org) (27) and the ff14SB force field (28). Using the Amber Tool LEaP, all systems were electrically neutralized with Na+ ions and solvated with TIP3P (29) water molecules. The receptor-binding domain complexed with ACE2 was solvated in a water box with the shape of a truncated octahedron and a distance of at least 25 Å from the borders to the solute.

The simulations followed a previously applied protocol (30). First, minimization was carried out in three consecutive steps to optimize the geometry of the initial structures. In the first step of the minimization, the water molecules were minimized, while all other atoms were restrained at the initial positions with a constant force of 10 kcal·mol^−1^·Å^−2^. In the second step, additional relaxation of the sodium ions and the hydrogen atoms of the protein was allowed, while the remaining protein was restrained with 10 kcal·mol^− 1^·Å^−2^. In the last step, no restraints were used, so the entire protein, ions, and water molecules were minimized. All three minimization parts started with 2500 steps using the steepest descent algorithm, followed by 2500 steps of a conjugate gradient minimization. After minimization, the systems were equilibrated in two successive steps. In the first step, the temperature was increased from 10 to 310 K within 0.1 ns and the protein was restrained with a constant force of 5 kcal·mol^−1^·Å^−2^. In the second step (0.4 ns length), only the Cα atoms of the protein were restrained with a constant force of 5 kcal·mol^−1^·Å^−2^. In both equilibration steps, the time step was 2 fs. Minimization and equilibration were carried out on CPUs, while the subsequent production runs were performed using pmemd.CUDA on Nvidia A100 GPUs (31–33). Subsequent 500 ns long production runs were performed without any restraints and at 310 K (regulated by a Berendsen thermostat (34)). Furthermore, the constant pressure periodic boundary conditions with an average pressure of 1 bar and isotropic position scaling were used. For bonds involving hydrogen, the SHAKE algorithm (35) was applied in the equilibration and production phases. To accelerate the production phase of the molecular dynamics (MD) simulations, hydrogen mass repartitioning (HMR) (36) was used in combination with a time step of 4 fs. For all different spike protein variants in complex with ACE2, the MD simulations were performed four times.

Trajectory analysis (analysis of root-mean-square fluctuations (RMSF), analysis of contacts (always with distance criterion of ≤5 Å between any pair of atoms; total fraction of contacts for residue pairs), measurement of interatomic distances, calculation of linear interaction energy (electrostatic and van der Waals interactions) was performed using the Amber tool cpptraj (37). For the calculation of interaction energies between ligand and receptor based on the MM/GBSA method, the mm_pbsa.pl script with default parameters was used (38–40).

### Statistics and display

Statistical analyses were performed with GraphPad Prism (version 8.0.0 for Windows, GraphPad Software, San Diego, California USA, www.graphpad.com) and statistical tests were applied as indicated below the figure. Plots were created in GraphPad and Gnuplot (version 5.2). All structure images were made with UCSF Chimera 1.15 (41).

## Acknowledgments

The authors gratefully acknowledge the compute resources and support provided by the Erlangen Regional Computing Center (RRZE) and by NHR@FAU.

## Author Contributions

E.S., F.Z., P.A. conceived the study; E.S. conducted the MD simulations; E.S., M.C., P.A. performed data analysis; E.S., L.H., P.A. generated visualization of the data; E.S., H.S., F.P., F.Z., P.A. contributed to the design of the study as well as discussion of the data; E. S., F.Z., P.A. wrote the initial draft; all authors reviewed the manuscript prior to submission.

---

**Supplemental Figure 1.**
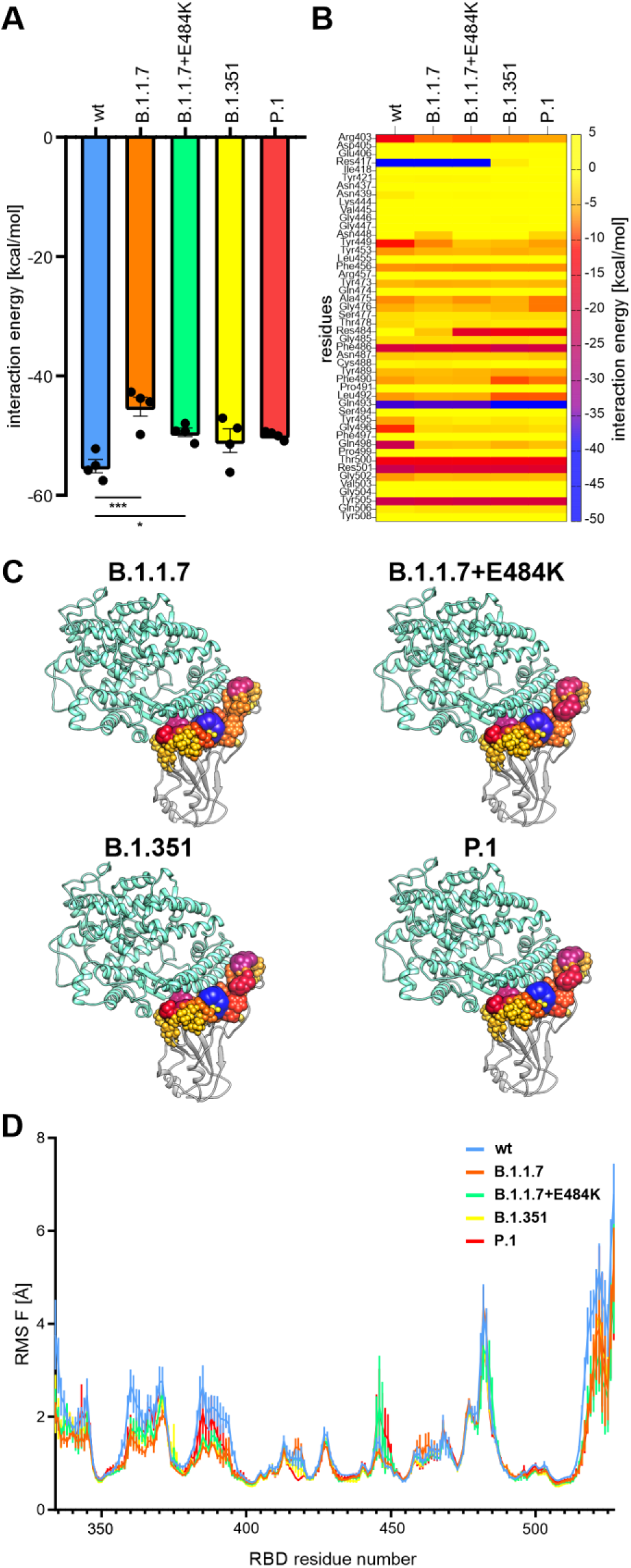
Decomposition of overall interaction energies and root mean square fluctuation (RMSF) values. (A) Interaction energies of the MM/GBSA analyses for the different receptor-binding domain (RBD) variants complexed with ACE2 (one-way ANOVA; * p<0.05, *** p<0.001). (B) Heat map for overall linear interaction energies for all residues of the RBD that were in a maximum distance of 8 Å with ACE2. Colors are according to the interaction energy from strong (blue) to weak (yellow). (C) Structural representation of RBD from different viral variants in complex with ACE2. The interacting residues within a maximum distance of 8 Å are shown according to their linear interaction energy in different colors and sphere diameters. (D) Root mean square fluctuation (RMSF) values for the different RBD variants show only subtle changes across different spike protein variants.

**Supplemental Figure 2.**
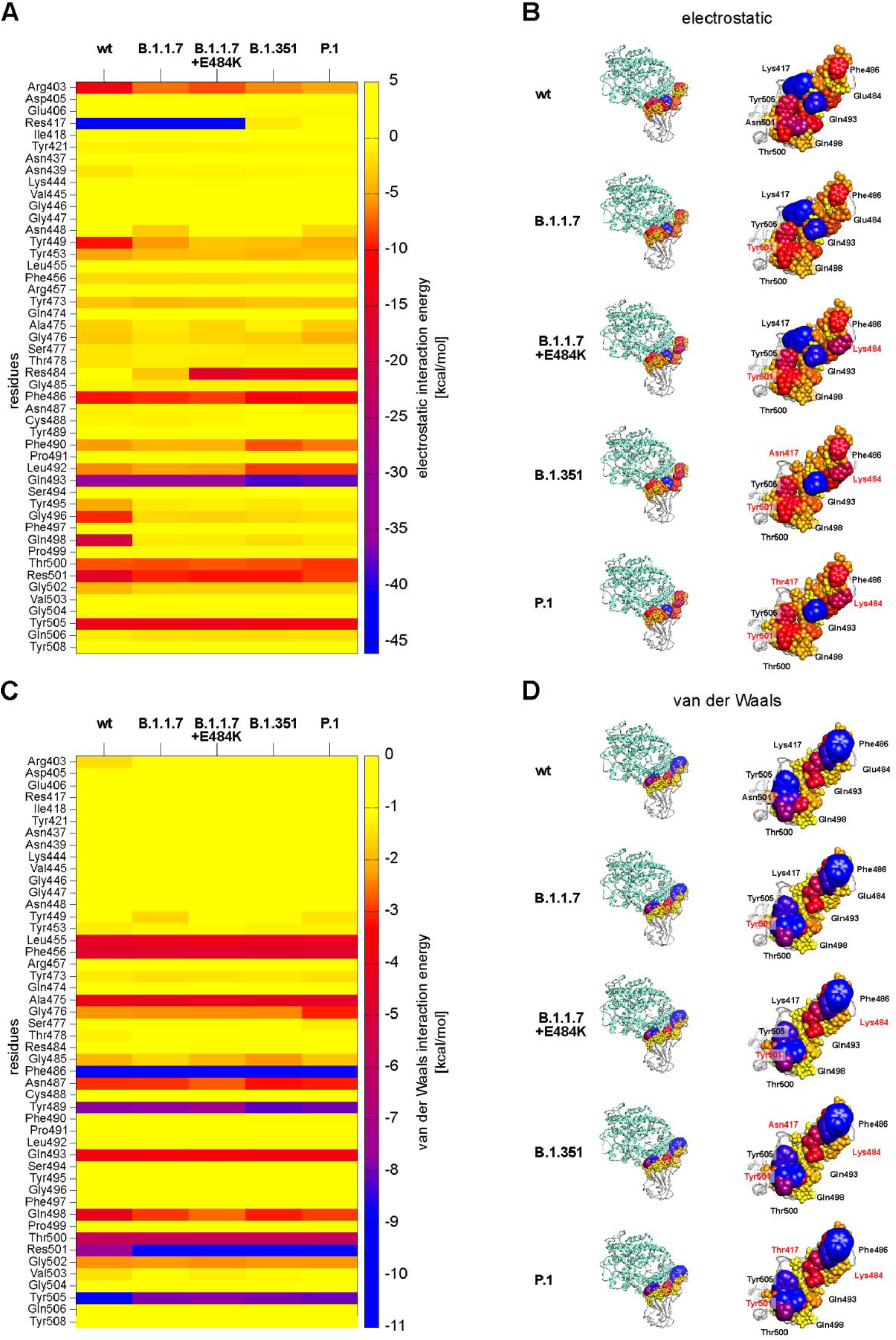
Decomposition into electrostatic and van der Waals linear interaction energies. (A) Heat map for electrostatic linear interaction energies for all residues of the receptor-binding domain (RBD) that were in a maximum distance of 8 Å with ACE2. Colors are according to the electrostatic linear interaction energy from strong (blue) to weak (yellow). (B) The different RBD variants are shown in complex with ACE2 (left) and residues are shown in different colors (blue-yellow) and spheres of different diameter according to their electrostatic linear interaction energy with ACE2. The residues interacting with ACE2 are also shown without ACE2 as a separate view (right). (C) Heat map for van der Waals linear interaction energies for all residues of the receptor-binding domain (RBD) that were in a maximum distance of 8 Å with ACE2. Colors are according to the van der Waals linear interaction energy from strong (blue) to weak (yellow). (D) The different RBD variants are shown in complex with ACE2 (left) and residues are shown in different colors (blue-yellow) and spheres of different diameters according to their van der Waals linear interaction energy with ACE2. The residues interacting with ACE2 are also shown without ACE2 as a separate view (right).

**Supplemental table.**

| Supplemental Table 1 |  |  |  |  |  |
| --- | --- | --- | --- | --- | --- |
| Number of families | 5 |  |  |  |  |
| Number of comparisons per family | 10 |  |  |  |  |
| Alpha | 0,05 |  |  |  |  |
| Tukey's multiple comparisons test | Mean Diff, | 95,00% CI of | Significant? | Summary | Adjusted P Value |
| Asp38 |  |  |  |  |  |
| wt vs. B.1.1.7 | 16,40 | 5,076 to 27,7 | Yes | * | 0,0179 |
| wt vs. B.1.1.7 +E484K | 16,50 | 5,157 to 27,8 | Yes | * | 0,0177 |
| wt vs. B.1.351 | 16,23 | 4,930 to 27,5 | Yes | * | 0,0182 |
| wt vs. P.1 | 16,48 | 5,136 to 27,8 | Yes | * | 0,0178 |
| B.1.1.7 vs. B.1.1.7 +E484K | 0,1000 | -0,3760 to 0,5 | No | ns | 0,8345 |
| B.1.1.7 vs. B.1.351 | -0,1750 | -0,8704 to 0,5 | No | ns | 0,8459 |
| B.1.1.7 vs. P.1 | 0,07500 | -0,3719 to 0,5 | No | ns | 0,9471 |
| B.1.1.7 +E484K vs. B.1.351 | -0,2750 | -1,037 to 0,4 | No | ns | 0,4793 |
| B.1.1.7 +E484K vs. P.1 | -0,02500 | -0,2618 to 0,2 | No | ns | 0,9889 |
| B.1.351 vs. P.1 | 0,2500 | -0,4729 to 0,9 | No | ns | 0,5647 |
| Tyr41 |  |  |  |  |  |
| wt vs. B.1.1.7 | 24,53 | 11,77 to 37,2 | Yes | ** | 0,0058 |
| wt vs. B.1.1.7 +E484K | 28,25 | 23,92 to 32,5 | Yes | **** | <0,0001 |
| wt vs. B.1.351 | 20,40 | 7,561 to 33,2 | Yes | * | 0,0112 |
| wt vs. P.1 | 23,68 | 8,978 to 38,3 | Yes | * | 0,0112 |
| B.1.1.7 vs. B.1.1.7 +E484K | 3,725 | -9,080 to 16,5 | No | ns | 0,6722 |
| B.1.1.7 vs. B.1.351 | -4,125 | -17,87 to 9,6 | No | ns | 0,7893 |
| B.1.1.7 vs. P.1 | -0,8500 | -15,61 to 13,9 | No | ns | 0,9993 |
| B.1.1.7 +E484K vs. B.1.351 | -7,850 | -20,74 to 5,0 | No | ns | 0,1942 |
| B.1.1.7 +E484K vs. P.1 | -4,575 | -19,32 to 10,2 | No | ns | 0,6159 |
| B.1.351 vs. P.1 | 3,275 | -11,52 to 18,0 | No | ns | 0,9102 |
| Gln42 |  |  |  |  |  |
| wt vs. B.1.1.7 | 6,575 | -7,610 to 20,7 | No | ns | 0,4456 |
| wt vs. B.1.1.7 +E484K | 9,375 | -6,492 to 25,2 | No | ns | 0,1840 |
| wt vs. B.1.351 | 5,000 | -9,151 to 19,1 | No | ns | 0,6494 |
| wt vs. P.1 | 7,125 | -7,013 to 21,2 | No | ns | 0,3536 |
| B.1.1.7 vs. B.1.1.7 +E484K | 2,800 | -7,675 to 13,2 | No | ns | 0,6623 |
| B.1.1.7 vs. B.1.351 | -1,575 | -11,92 to 8,7 | No | ns | 0,9747 |
| B.1.1.7 vs. P.1 | 0,5500 | -9,166 to 10,2 | No | ns | 0,9994 |
| B.1.1.7 +E484K vs. B.1.351 | -4,375 | -14,49 to 5,7 | No | ns | 0,3449 |
| B.1.1.7 +E484K vs. P.1 | -2,250 | -10,75 to 6,2 | No | ns | 0,6699 |
| B.1.351 vs. P.1 | 2,125 | -7,356 to 11,6 | No | ns | 0,9052 |
| Leu45 |  |  |  |  |  |
| wt vs. B.1.1.7 | -5,875 | -12,72 to 0,9 | No | ns | 0,0812 |
| wt vs. B.1.1.7 +E484K | -3,675 | -13,22 to 5,8 | No | ns | 0,6016 |
| wt vs. B.1.351 | -7,025 | -14,13 to 0,0 | No | ns | 0,0518 |
| wt vs. P.1 | -5,500 | -19,31 to 8,3 | No | ns | 0,5153 |
| B.1.1.7 vs. B.1.1.7 +E484K | 2,200 | -7,739 to 12,1 | No | ns | 0,8211 |
| B.1.1.7 vs. B.1.351 | -1,150 | -4,006 to 1,7 | No | ns | 0,5615 |
| B.1.1.7 vs. P.1 | 0,3750 | -14,53 to 15,2 | No | ns | >0,9999 |
| B.1.1.7 +E484K vs. B.1.351 | -3,350 | -13,59 to 6,8 | No | ns | 0,5576 |
| B.1.1.7 +E484K vs. P.1 | -1,825 | -15,65 to 12,0 | No | ns | 0,9820 |
| B.1.351 vs. P.1 | 1,525 | -13,64 to 16,6 | No | ns | 0,9794 |
| Lys353 |  |  |  |  |  |
| wt vs. B.1.1.7 | 49,18 | 30,38 to 67,9 | Yes | ** | 0,0030 |
| wt vs. B.1.1.7 +E484K | 49,38 | 30,32 to 68,4 | Yes | ** | 0,0033 |
| wt vs. B.1.351 | 49,78 | 30,73 to 68,8 | Yes | ** | 0,0032 |
| wt vs. P.1 | 49,95 | 30,93 to 68,9 | Yes | ** | 0,0031 |
| B.1.1.7 vs. B.1.1.7 +E484K | 0,2000 | -2,148 to 2,5 | No | ns | 0,9901 |
| B.1.1.7 vs. B.1.351 | 0,6000 | -1,724 to 2,9 | No | ns | 0,7259 |
| B.1.1.7 vs. P.1 | 0,7750 | -1,446 to 2,9 | No | ns | 0,5814 |
| B.1.1.7 +E484K vs. B.1.351 | 0,4000 | -0,1769 to 0,9 | No | ns | 0,1840 |
| B.1.1.7 +E484K vs. P.1 | 0,5750 | -0,2943 to 1,4 | No | ns | 0,1858 |
| B.1.351 vs. P.1 | 0,1750 | -0,6945 to 1,0 | No | ns | 0,9186 |

## References

1. N. Chen, M. Zhou, X. Dong, J. Qu, F. Gong, Y. Han, Y. Qiu, J. Wang, Y. Liu, Y. Wei, J. Xia, T. Yu, X. Zhang, L. Zhang, Epidemiological and clinical characteristics of 99 cases of 2019 novel coronavirus pneumonia in Wuhan, China: a descriptive study. The Lancet 395, 507–513 (2020).

2. World Health Organization, WHO Coronavirus (COVID-19) Dashboard. https://covid19.who.int.

3. M. Hoffmann, H. Kleine-Weber, S. Schroeder, N. Krüger, T. Herrler, S. Erichsen, T. S. Schiergens, G. Herrler, N.-H. Wu, A. Nitsche, M. A. Müller, C. Drosten, S. Pöhlmann, SARS-CoV-2 Cell Entry Depends on ACE2 and TMPRSS2 and Is Blocked by a Clinically Proven Protease Inhibitor. Cell 181, 271–280.e8 (2020).

4. P. Zhou, X.-L. Yang, X.-G. Wang, B. Hu, L. Zhang, W. Zhang, H.-R. Si, Y. Zhu, B. Li, C.-L. Huang, H.-D. Chen, J. Chen, Y. Luo, H. Guo, R.-D. Jiang, M.-Q. Liu, Y. Chen, X.-R. Shen, X. Wang, X.-S. Zheng, K. Zhao, Q.-J. Chen, F. Deng, L.-L. Liu, B. Yan, F.-X. Zhan, Y.-Y. Wang, G.-F. Xiao, Z.-L. Shi, A pneumonia outbreak associated with a new coronavirus of probable bat origin. Nature 579, 270–273 (2020).

5. J. Yang, S. J. L. Petitjean, M. Koehler, Q. Zhang, A. C. Dumitru, W. Chen, S. Derclaye, S. P. Vincent, P. Soumillion, D. Alsteens, Molecular interaction and inhibition of SARS-CoV-2 binding to the ACE2 receptor. Nature communications 11, 4541 (2020).

6. W. Cao, C. Dong, S. Kim, D. Hou, W. Tai, L. Du, W. Im, X. F. Zhang, Biomechanical characterization of SARS-CoV-2 spike RBD and human ACE2 protein-protein interaction. Biophysical journal. 10.1016/j.bpj.2021.02.007 (2021).

7. M. Hoffmann, H. Kleine-Weber, S. Pöhlmann, A Multibasic Cleavage Site in the Spike Protein of SARS-CoV-2 Is Essential for Infection of Human Lung Cells. Molecular cell 78, 779–784.e5 (2020).

8. G. Simmons, P. Zmora, S. Gierer, A. Heurich, S. Pöhlmann, Proteolytic activation of the SARS-coronavirus spike protein: cutting enzymes at the cutting edge of antiviral research. Antiviral research 100, 605–614 (2013).

9. S. Belouzard, V. C. Chu, G. R. Whittaker, Activation of the SARS coronavirus spike protein via sequential proteolytic cleavage at two distinct sites. Proceedings of the National Academy of Sciences of the United States of America 106, 5871–5876 (2009).

10. K. K. Chan, D. Dorosky, P. Sharma, S. A. Abbasi, J. M. Dye, D. M. Kranz, A. S. Herbert, E. Procko, Engineering human ACE2 to optimize binding to the spike protein of SARS coronavirus 2. Science (New York, N.Y.) 369, 1261–1265 (2020).

11. J. K. Millet, G. R. Whittaker, Host cell entry of Middle East respiratory syndrome coronavirus after two-step, furin-mediated activation of the spike protein. Proceedings of the National Academy of Sciences of the United States of America 111, 15214–15219 (2014).

12. S. Matsuyama, M. Ujike, S. Morikawa, M. Tashiro, F. Taguchi, Protease-mediated enhancement of severe acute respiratory syndrome coronavirus infection. Proceedings of the National Academy of Sciences of the United States of America 102, 12543–12547 (2005).

13. D. Ni, K. Lau, F. Lehmann, A. Fränkl, D. Hacker, F. Pojer, H. Stahlberg, Structural investigation of ACE2 dependent disassembly of the trimeric SARS-CoV-2 Spike glycoprotein. 10.1101/2020.10.12.336016 (2020).

14. D. J. Benton, A. G. Wrobel, P. Xu, C. Roustan, S. R. Martin, P. B. Rosenthal, J. J. Skehel, S. J. Gamblin, Receptor binding and priming of the spike protein of SARS-CoV-2 for membrane fusion. Nature 588, 327–330 (2020).

15. Centers for Disease Control and Prevention, Science Brief: Emerging SARS-CoV-2 Variants. https://www.cdc.gov/coronavirus/2019-ncov/more/science-and-research/scientific-brief-emerging-variants.html.

16. World Health Organization, Weekly epidemiological update - 2 February 2021. https://www.who.int/publications/m/item/weekly-epidemiological-update2-february-2021.

17. S. Jangra, C. Ye, R. Rathnasinghe, D. Stadlbauer, F. Krammer, V. Simon, L. Martinez-Sobrido, A. Garcia-Sastre, M. Schotsaert, The E484K mutation in the SARS-CoV-2 spike protein reduces but does not abolish neutralizing activity of human convalescent and post-vaccination sera. medRxiv: the preprint server for health sciences. 10.1101/2021.01.26.21250543 (2021).

18. Y. Weisblum, F. Schmidt, F. Zhang, J. DaSilva, D. Poston, J. C. Lorenzi, F. Muecksch, M. Rutkowska, H.-H. Hoffmann, E. Michailidis, C. Gaebler, M. Agudelo, A. Cho, Z. Wang, A. Gazumyan, M. Cipolla, L. Luchsinger, C. D. Hillyer, M. Caskey, D. F. Robbiani, C. M. Rice, M. C. Nussenzweig, T. Hatziioannou, P. D. Bieniasz, Escape from neutralizing antibodies by SARS-CoV-2 spike protein variants. eLife 9 (2020).

19. T. Zhou, Y. Tsybovsky, J. Gorman, M. Rapp, G. Cerutti, G.-Y. Chuang, P. S. Katsamba, J. M. Sampson, A. Schön, J. Bimela, J. C. Boyington, A. Nazzari, A. S. Olia, W. Shi, M. Sastry, T. Stephens, J. Stuckey, I.-T. Teng, P. Wang, S. Wang, B. Zhang, R. A. Friesner, D. D. Ho, J. R. Mascola, L. Shapiro, P. D. Kwong, Cryo-EM Structures of SARS-CoV-2 Spike without and with ACE2 Reveal a pH-Dependent Switch to Mediate Endosomal Positioning of Receptor-Binding Domains. Cell host & microbe 28, 867–879.e5 (2020).

20. R. P. Walensky, H. T. Walke, A. S. Fauci, SARS-CoV-2 Variants of Concern in the United States-Challenges and Opportunities. JAMA. 10.1001/jama.2021.2294 (2021).

21. S. Chakraborty, Evolutionary and structural analysis elucidates mutations on SARS-CoV2 spike protein with altered human ACE2 binding affinity. Biochemical and biophysical research communications 534, 374–380 (2021).

22. M. H. Cheng, J. M. Krieger, B. Kaynak, M. Arditi, I. Bahar, Impact of South African 501.V2 Variant on SARS-CoV-2 Spike Infectivity and Neutralization: A Structure-based Computational Assessment. 10.1101/2021.01.10.426143 (2021).

23. T. N. Starr, A. J. Greaney, S. K. Hilton, D. Ellis, K. H. D. Crawford, A. S. Dingens, M. J. Navarro, J. E. Bowen, M. A. Tortorici, A. C. Walls, N. P. King, D. Veesler, J. D. Bloom, Deep Mutational Scanning of SARS-CoV-2 Receptor Binding Domain Reveals Constraints on Folding and ACE2 Binding. Cell 182, 1295–1310.e20 (2020).

24. C. K. Wibmer, F. Ayres, T. Hermanus, M. Madzivhandila, P. Kgagudi, B. E. Lambson, M. Vermeulen, K. van den Berg, T. Rossouw, M. Boswell, V. Ueckermann, S. Meiring, A. von Gottberg, C. Cohen, L. Morris, J. N. Bhiman, P. L. Moore, SARS-CoV-2 501Y.V2 escapes neutralization by South African COVID-19 donor plasma. bioRxiv: the preprint server for biology. 10.1101/2021.01.18.427166 (2021).

25. B. Qiao, M. La Olvera de Cruz, Enhanced Binding of SARS-CoV-2 Spike Protein to Receptor by Distal Polybasic Cleavage Sites. ACS nano 14, 10616–10623 (2020).

26. B. Dehury, V. Raina, N. Misra, M. Suar, Effect of mutation on structure, function and dynamics of receptor binding domain of human SARS-CoV-2 with host cell receptor ACE2: a molecular dynamics simulations study. Journal of biomolecular structure & dynamics. 10.1080/07391102.2020.1802348, 1–15 (2020).

27. D. A. Case, K. Belfon, I. Y. Ben-Shalom, S. R. Brozell, D. S. Cerutti, T. E. Cheatham, III, V. W. D. Cruzeiro, T. A. Darden, R. E. Duke, G. Giambasu, M. K. Gilson, H. Gohlke, A. W. Goetz, R. Harris, S. Izadi, S. A. Izmailov, K. Kasavajhala, A. Kovalenko, R. Krasny, T. Kurtzman, T. S. Lee, S. LeGrand, P. Li, C. Lin, J. Liu, T. Luchko, R. Luo, V. Man, K. M. Merz, Y. Miao, O. Mikhailovskii, G. Monard, H. Nguyen, A. Onufriev, F. Pan, S. Pantano, R. Qi, D. R. Roe, A. Roitberg, C. Sagui, S. Schott-Verdugo, J. Shen, C. L. Simmerling, N. R. Skrynnikov, J. Smith, J. Swails, R. C. Walker, J. Wang, L. Wilson, R. M. Wolf, X. Wu, Y. Xiong, Y. Xue, D. M. York, P. A. Kollman, AMBER 2020.

28. J. A. Maier, C. Martinez, K. Kasavajhala, L. Wickstrom, K. E. Hauser, C. Simmerling, ff14SB: Improving the Accuracy of Protein Side Chain and Backbone Parameters from ff99SB. Journal of chemical theory and computation 11, 3696–3713 (2015).

29. W. L. Jorgensen, J. Chandrasekhar, J. D. Madura, R. W. Impey, M. L. Klein, Comparison of simple potential functions for simulating liquid water. The Journal of Chemical Physics 79, 926–935 (1983).

30. E. Socher, H. Sticht, A. H. C. Horn, The conformational stability of nonfibrillar amyloid-β peptide oligomers critically depends on the C-terminal peptide length. ACS chemical neuroscience 5, 161–167 (2014).

31. R. Salomon-Ferrer, A. W. Götz, D. Poole, S. Le Grand, R. C. Walker, Routine Microsecond Molecular Dynamics Simulations with AMBER on GPUs. 2. Explicit Solvent Particle Mesh Ewald. Journal of chemical theory and computation 9, 3878–3888 (2013).

32. A. W. Götz, M. J. Williamson, D. Xu, D. Poole, S. Le Grand, R. C. Walker, Routine Microsecond Molecular Dynamics Simulations with AMBER on GPUs. 1. Generalized Born. Journal of chemical theory and computation 8, 1542–1555 (2012).

33. S. Le Grand, A. W. Götz, R. C. Walker, SPFP: Speed without compromise—A mixed precision model for GPU accelerated molecular dynamics simulations. Computer Physics Communications 184, 374–380 (2013).

34. H. J. C. Berendsen, J. P. M. Postma, W. F. van Gunsteren, A. DiNola, J. R. Haak, Molecular dynamics with coupling to an external bath. The Journal of Chemical Physics 81, 3684–3690 (1984).

35. J.-P. Ryckaert, G. Ciccotti, H. J. Berendsen, Numerical integration of the cartesian equations of motion of a system with constraints: molecular dynamics of n-alkanes. Journal of Computational Physics 23, 327–341 (1977).

36. C. W. Hopkins, S. Le Grand, R. C. Walker, A. E. Roitberg, Long-Time-Step Molecular Dynamics through Hydrogen Mass Repartitioning. Journal of chemical theory and computation 11, 1864–1874 (2015).

37. D. R. Roe, T. E. Cheatham, PTRAJ and CPPTRAJ: Software for Processing and Analysis of Molecular Dynamics Trajectory Data. Journal of chemical theory and computation 9, 3084–3095 (2013).

38. S. Genheden, U. Ryde, The MM/PBSA and MM/GBSA methods to estimate ligand-binding affinities. Expert opinion on drug discovery 10, 449–461 (2015).

39. P. A. Kollman, I. Massova, C. Reyes, B. Kuhn, S. Huo, L. Chong, M. Lee, T. Lee, Y. Duan, W. Wang, O. Donini, P. Cieplak, J. Srinivasan, D. A. Case, T. E. Cheatham, Calculating structures and free energies of complex molecules: combining molecular mechanics and continuum models. Accounts of chemical research 33, 889–897 (2000).

40. W. Wang, O. Donini, C. M. Reyes, P. A. Kollman, Biomolecular simulations: recent developments in force fields, simulations of enzyme catalysis, protein-ligand, protein-protein, and protein-nucleic acid noncovalent interactions. Annual review of biophysics and biomolecular structure 30, 211–243 (2001).

41. E. F. Pettersen, T. D. Goddard, C. C. Huang, G. S. Couch, D. M. Greenblatt, E. C. Meng, T. E. Ferrin, UCSF Chimera--a visualization system for exploratory research and analysis. Journal of computational chemistry 25, 1605–1612 (2004).

